# Mapping the evolutionary path towards multi-drug resistance in the pandemic *Escherichia coli* ST131 lineage

**DOI:** 10.64898/2025.12.08.692488

**Authors:** Anna K. Pöntinen, Nguyen Vinh Trung, Sudaraka Mallawaarachchi, Rebecca A. Gladstone, Juri Kuronen, Ørjan Samuelsen, Huynh Xuan Yen, Phung Le Kim Yen, Nguyen Phu Huong Lan, Nguyen Thanh Dung, Nguyen Van Vinh Chau, Julian Parkhill, Gerry Tonkin-Hill, Ngo Thi Hoa, Jukka Corander

**Author notes:** Corresponding authors: Anna K. Pöntinen and Jukka Corander.

## Abstract

*Escherichia coli* sequence type (ST) 131 is the most widely studied genetic lineage of the species to date, originally identified in the early 2000s as an increasingly common cause of human urinary tract and bloodstream infections worldwide. This lineage is subdivided into four extant main subclades A, B, C1 and C2 that exhibit distinct features in terms of invasiveness, antibiotic resistance and transmissibility. However, the evolutionary pathway from the generally susceptible ST131-B to the drug-resistant ST131-C clades remains poorly mapped. To fill this knowledge gap, we analysed in detail human clinical isolates obtained in Vietnam, designated as belonging to the generally neglected minor clade ST131-B0. We sequenced them using both short- and long-read technology, and combined these data with a recently published high-resolution genomic collection to provide further insight into the evolutionary process and its timeline. Extensive genomic analyses established ST131-B0 as an intermediary progenitor in the evolutionary path leading from ST131-B to the ST131-C clades, associated with multiple type I pili switches, as well as the loss and gain of specific chromosomal genes representing diverse core functions such as metabolism, transcription, DNA binding and type II toxin-antitoxin systems. Furthermore, all Vietnamese isolates of ST131-B0 unprecedentedly harboured *bla*_CTX-M_ genes encoding extended-spectrum β-lactamases, a trait dominant in ST131-C clades and not previously observed in ST131-B0. Our study supports the hypothesis that the ST131-C clades have gradually evolved from ST131-B by reducing the host range with better adaptation to colonising humans under selective conditions.

## Introduction

*Escherichia coli* is a leading causative agent of human bloodstream infections (BSI) and urinary tract infections (UTI) globally with high associated morbidity and mortality (Foxman 2014; Kern and Rieg 2020; Bonten et al. 2021). The clinical concern is exacerbated by the high prevalence of multi-drug resistant (MDR) *E. coli*, reducing therapeutic options and, thereby, prolonging hospitalisation and increasing mortality. In particular, the increase in the prevalence of extended-spectrum β-lactamase (ESBL)-producing *E. coli* remains a global concern (Naylor et al. 2019; Shamsrizi et al. 2020; WHO 2024). Among extraintestinal pathogenic *E. coli* (ExPEC), sequence type (ST) 131 has generally held a predominant position worldwide among ESBL-producers since the early 2000s (Nicolas-Chanoine et al. 2007; Nicolas-Chanoine et al. 2014; Kallonen et al. 2017; Gladstone et al. 2021). Based on representative genomic surveillance, the prevalence of ST131 rapidly increased, such that it became one of the leading causes of invasive ExPEC infections, followed by a new equilibrium within the ExPEC population (Kallonen et al. 2017; Gladstone et al. 2021; Pöntinen et al. 2024).

Based on representative longitudinal sampling and modelling, antimicrobial resistance (AMR) appears not to be the sole driver of the epidemiological success of ST131 (Gladstone et al. 2021; Pöntinen et al. 2024). It has been suggested that a combination of negative frequency-dependent selection (NFDS) within a wider commensal niche and particular selection pressures deriving from non-penicillin β-lactam usage have resulted in differing equilibrium prevalences of ST131 in separate geographical regions (Kallonen et al. 2017; McNally et al. 2019; Gladstone et al. 2021; Pöntinen et al. 2024). Furthermore, the sublineages of ST131, here referred to as ST131 clades A, B, C1, and C2, have been characterised by notably distinct associations with AMR traits. Of note, ST131-C1 and -C2 have been described as frequently carrying CTX-M-type ESBLs (Price et al. 2013; Petty et al. 2014; Brodrick et al. 2017; Gladstone et al. 2021). Typically, ST131-C2 has been associated with CTX-M-15, while ST131-C1 more commonly carries CTX-M-14 (Nicolas-Chanoine et al. 2014; Bevan et al. 2017; Komori et al. 2024). While found integrated into the chromosome (Coque et al. 2008; Hirai et al. 2013; Hamamoto et al. 2016), ESBL-encoding genes are also frequently carried on plasmids in *E. coli* (Cave et al. 2023; Arredondo-Alonso et al. 2025). Therefore, concern caused by increasing AMR levels is further exacerbated by the global dissemination of readily mobile plasmid-borne AMR determinants. Furthermore, a study by Molari et al. (2025) indicated a broader role of the accessory genes in *E. coli* evolution by extensive accessory genome variation within *E. coli* ST131, as well as structural modifications at rates comparable to individual nucleotide substitutions (Molari et al. 2025).

Human colonisation represents the main reservoir for ExPECs, and ST131 is, accordingly, capable of asymptomatically residing in the human intestines (Sarkar et al. 2018; Johnson et al. 2022). However, ST131-B, in particular, has been shown to represent a zoonotic pathogen (Platell et al. 2011; Liu et al. 2018; Reid et al. 2019). Animal ST131-B isolates are particularly associated with poultry (Saidenberg et al. 2020), and the contaminated products thereof are proposed as potential foodborne vehicles for human infections caused by *E. coli* ST131 (Manges 2016; Liu et al. 2018; Roer et al. 2019). Although generally reported as more susceptible to antimicrobials compared to the other ST131 clades, ST131-B (ST131-*H*22) isolates with both human BSI and poultry origin have been reported to sporadically carry plasmid-mediated colistin resistance (*mcr*) traits (Roer et al. 2019; Saidenberg et al. 2020). However, in a study of uropathogenic *E. coli* infections, the clinical presentation within the human urinary tract did not differ significantly between ST131-B and ST131-C, as the clades were equally likely manifested as asymptomatic bacteriuria (Liu et al. 2018). Furthermore, an investigation of the gut colonisation in neonates and the relative invasiveness of *E. coli* lineages showed that ST131-B was more associated with asymptomatic human colonisation compared with ST131-C clades (Mäklin et al. 2022; Pöntinen et al. 2024). Combined, these findings suggest that ST131-B has adapted to two different primary niches, one representing asymptomatic circulation among humans and the other colonisation of poultry, both of which then contribute towards the human infection burden. The genetic determinants behind these adaptations are potentially partly overlapping, but not currently fully resolved, see the discussion in Arredondo-Alonso et al. (Arredondo-Alonso et al. 2025). Finally, it is worth noticing that the estimated transmissibility for ST131-C1 and ST131-C2 is significantly lower compared with ST131-A (sufficient data not available for ST131-B), suggesting that ST131-C clades may be more adept at transmission within the healthcare settings aided by antibiotic selection pressure than in community settings (Ojala et al. 2025).

The current evidence suggests that a branch of ST131-B has over time evolved to the minor subclade ST131-C0 and further to ST131-C1 and ST131-C2 clades by gradual *fimH* transition and by acquiring fluoroquinolone resistance mutations and plasmid-borne *bla*_CTX-M_ genes in the ST131-C clades (Petty et al. 2014; Ben Zakour et al. 2016; Matsumura et al. 2016; Stoesser et al. 2016; Pitout and Finn 2020). However, due to the limited number of intermediary isolates, the full evolutionary path from ST131-B ancestor to ST131-C clades has remained elusive. Here, we introduce a collection of clinical ST131-B0 isolates, sequenced by using both short- and long-read techniques, and combine these data with ST131 genomes from a recent high-resolution sequenced *E. coli* BSI cohort (Gladstone et al. 2021; Arredondo-Alonso et al. 2025) to fill this prevailing knowledge gap. We show that ST131-B0 unprecedentedly carries CTX-M-type ESBLs both on plasmids and chromosomes and, combined with high-resolution dating analyses in context, we map in detail how ST131-B0 arose from ST131-B, and further evolved into the present predominantly MDR pathogen lineage ST131-C.

## Results

### Isolate collection and population structure

In total, 212 *E. coli* ST131-B and -C clinical isolates were analysed within the study (Supp. Table 1, Microreact project [https://microreact.org/project/ecoli-st131-evolution]), of which 9 were from the Oxford University Clinical Research Unit (OUCRU), Vietnam (sequenced in this study) and 203 from the Norwegian surveillance programme on resistant microbes (NORM), Norway (Gladstone et al. 2021). Of note, a total of 206 hybrid assemblies was available for the combined collection. All OUCRU isolates belonged to the clade ST131-B0, while the NORM isolates were distributed between ST131-B (n=74), ST131-B0 (n=4), ST131-C0 (n=4), ST131-C1 (n=70), and ST131-C2 (n=51). Aligning the *fimH*-encoded type I pili with the dated phylogeny identified an apparent switch in the predominant *fimH* type per clade, from predominantly *fimH22* in ST131-B to *fimH30* in the ST131-C1 and ST131-C2, while *fimH54* was the most common type in ST131-B0 (Fig. 1). When screening for flagellar H and polysaccharide O antigens, all isolates were of H4 and, apart from one O2 in ST131-B, of O25 serotype (Fig. 1). The most abundant capsule type was K5, covering 52.8% (112/212) of the isolates across all ST131 clades. It has previously been found to be the ancestral capsule type in ST131-B and -C clades (Gladstone et al. 2025). Of note, 8 out of 9 of the OUCRU ST131-B0 isolates carried the K5 capsule, supporting their position as an intermediary progenitor to ST131-C where substantial capsule diversity has emerged later (Gladstone et al. 2025). The second most common capsule type overall was K2, covering 10.8% (23/212) of the total isolates and disseminated in all of the ST131 clades apart from ST131-C1. K100 was found in 4.7% (10/212) of the total isolates and exclusively in ST131-C2, and K16 in 3.8% (8/212), only in ST131-B. K1 type was found in 4 ST131-B and K20 type in one ST131-C2 isolate (Fig. 1). 25.5% (54/212) isolates remained with untypeable or unknown capsule loci.

**Figure 1.**
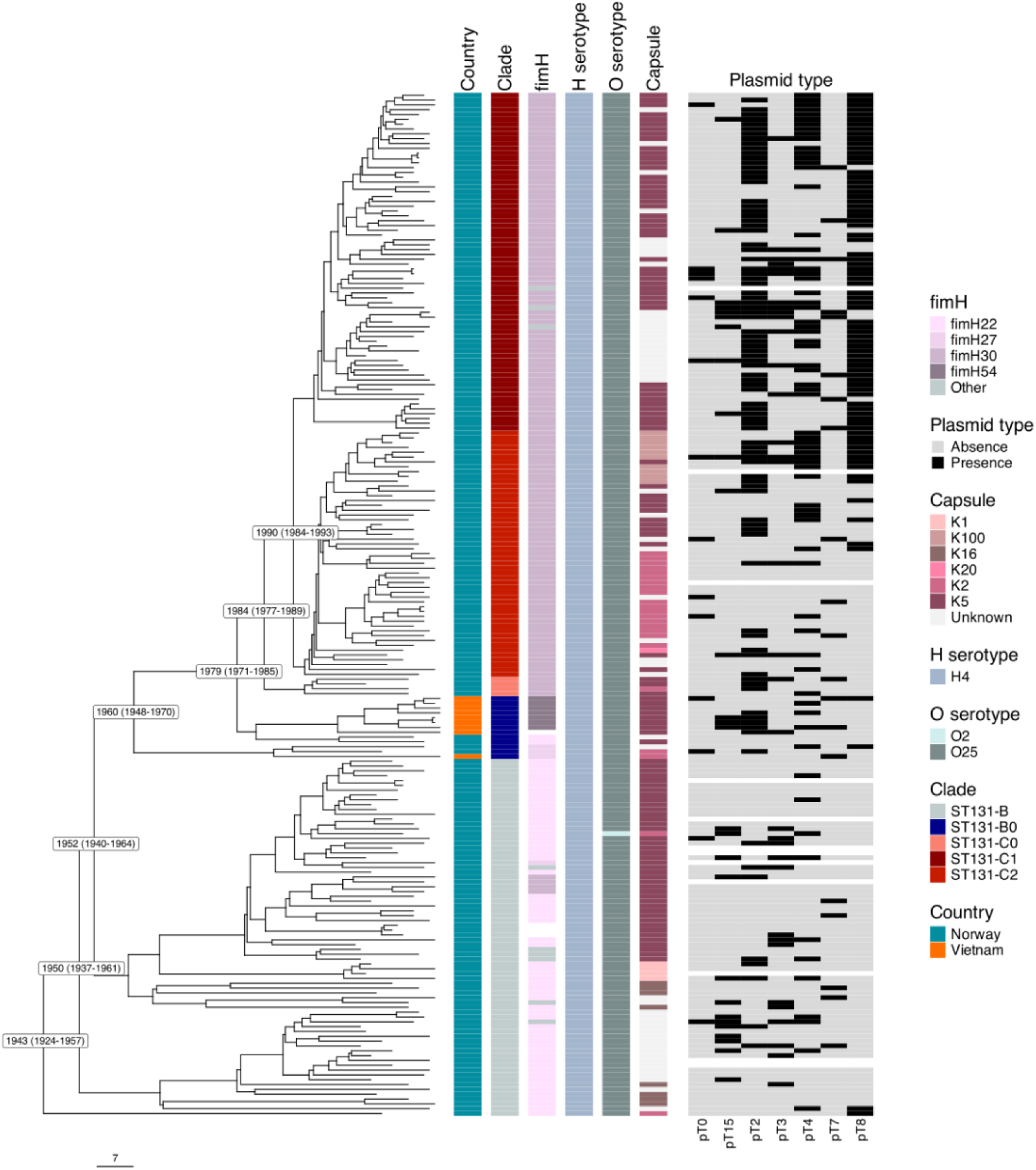
Dated phylogeny of the Vietnamese ST131-B0 (n=9) and Norwegian ST131-B (n=74), ST131-B0 (n=4), ST131-C0 (n=4), ST131-C1 (n=70) and ST131-C2 (n=51), aligned with *fimH*/O/H/K typing and presence/absence patterns of the selected plasmid types (pT) found in the Vietnamese ST131-B0, as indicated in the legend. Isolates lacking any plasmid-predicted contigs are shown as empty in the plasmid type panel. The tree scale indicates years. Time of the most recent common ancestor of each clade with 95% highest posterior density intervals is indicated within the inferred dated phylogeny.

### Evolutionary timeline from ST131-B to ST131-C

Dating analyses further support ST131-B0 as an intermediary progenitor of the ST131-C clades in the evolutionary path deriving from ST131-B (Fig. 1). The most recent common ancestor (MRCA) for the combined OUCRU and NORM ST131-B and ST131-C clades was estimated to date back to 1943 (95% highest posterior density [HPD] interval 1924-1957). ST131-B0 and ST131-C clades were estimated to diverge from the ancestral ST131-B clade in 1952 (95% HPD 1940-1964). Shortly after, estimated in 1960 (95% HPD 1958-1970), an older branch of ST131-B0 clade diverged from the rest of the clade, creating a deep split between the two branches. Finally, ST131-C clades were estimated to diverge from the more recent branch of ST131-B0 approximately two decades later in 1979 (95% HPD 1971-1985). Splits in the ST131-C clades were seen in more rapid succession, with ST131-C0 estimated to diverge in 1984 (95% HPD 1977-1989), and ST131-C1 and ST131-C2 clades in 1990 (95% HPD 1984-1993) (Fig. 1).

### CTX-M ESBL association with ST131 clades

In total, 6 different CTX-M type ESBLs were found in the combined collection: CTX-M-14, CTX-M-15, CTX-M-19, CTX-M-24, CTX-M-27 and CTX-M-127 (Fig. 2). The most common type was CTX-M-15, covering 20.9% (43/206) of the total isolates (hybrid assemblies), and was found in the OUCRU ST131-B0 and NORM ST131-C1 and ST131-C2. It was found equally often carried on plasmids (21/43) and chromosomes (22/43), and 6 of these isolates also carried more than one copy of the gene on their chromosomes. However, those located in chromosomes were most commonly (19/22) found in ST131-C2 (also 1 in ST131-C1 and 3 in OUCRU ST131-B0). CTX-M-27 was found in 9.7% (20/206) of the isolates, in ST131-C1 (15/20) and the OUCRU ST131-B0 (5/20). It was mostly plasmid-borne but 3 out of 20 isolates carried it on chromosomes. CTX-M-14 was less common and found only in 5 ST131-C1 isolates, carried on both chromosomes and plasmids. CTX-M-19 and −127 were found only in one ST131-C1 plasmid and ST131-C2 chromosome, respectively, while two copies of CTX-M-24 were found in a single ST131-C1 chromosome.

**Figure 2.**
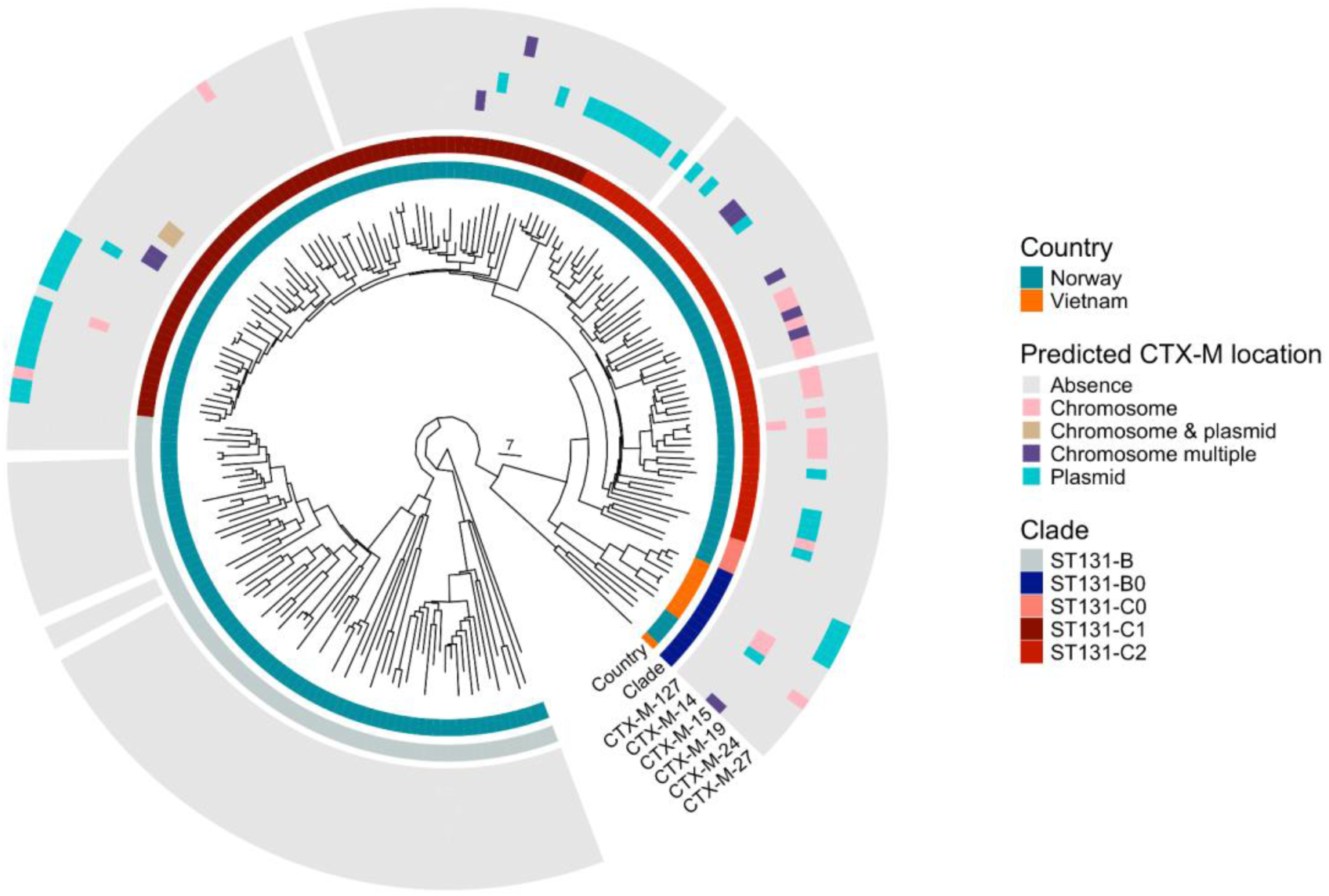
Predicted plasmid and chromosome locations of the CTX-M-type ESBLs in the Vietnamese and Norwegian ST131 isolates, aligned with the dated phylogeny where the tree scale indicates years. Presence of the *bla*_CTX-M_ genes is indicated by chromosome/plasmid affinity and by CTX-M type as colour coded in the legend. Isolates lacking hybrid assemblies are shown as empty.

### Comparative pangenome analysis and genome-wide co-selection

Of the 10,781 total genes within the combined ST131 collections (206 hybrid assemblies), 3964 (36.8%) were estimated as core genes (present in ≥99% of isolates), amounting to 80.7% (3964/4909) of the average genome size. Comparison of gene gain and loss rates between clades revealed the largest estimate of gene exchange relative to core branch length in ST131-C2 (Supp. Fig. 1A), statistically significantly different from both ST131-B0 (p=0.0455) and ST131-B (p=0.0049). Gene exchange rates of ST131-C1 were also significantly higher than those of ST131-B (p=0.0392). These differences were also seen in the cumulative gene gain and loss events (Supp. Fig. 1B).

Comparison of gene presence and absence between ST131-B0 and ST131-C clades (C1, C2) revealed specific genes and potential homologues gained and lost in the transition between these clades (Supp. Table 2). Of note, pangenomes were estimated with a minimum pairwise sequence-identity threshold of 70% between genes, and additional homologous genes might exist at lower sequence identities. In total, six unique genes were only present in ST131-B0. These were all predicted to be located on the chromosome and included genes functioning in arginine (*arcA*), ornithine (*argF*) and carbamate (*arcC*) metabolism and encoding a putative membrane transporter (*yfcC*). The eight unique genes gained by the ST131-C clades (absent in ST131-B0), included genes encoding oxidoreductase and transcription repressor (*bdcAR*), type II toxin-antitoxin system (*chpBS*), DNA-binding proteins, endoribonuclease (*yjgH*) and membrane protein (*yjgN*). These were also predominantly chromosomal as at least 95.9% (117/122) of the isolates harboured them on chromosome-predicted contigs, supporting the importance of both the lost and gained genes as essential to the core functionality of their respective subclades.

Investigation of potential co-selection and pangenome-spanning epistasis of genomic variation by LDWeaver (Mallawaarachchi et al. 2024) revealed several short- and long-range links between genomic variation, illustrated in network plots (Supp. Fig. 2). The top-ranking region among short-range linkage of variation with most significant links was found to be a site in *sitD*, encoding an iron/manganese ABC transporter permease subunit (Supp. Fig. 2A). The network analysis indicated potential co-selection of *sitD* amongst others with *ydaT*, encoding a bacterial toxin, and *bet,* encoding a phage recombination protein. The highest number of links was found between *sitD* and replication protein P (DOHBNP_06365). The top-ranking long-range linkage of variation revealed a complex network of multiple linked loci associated especially with metabolic processes (Supp. Fig. 2B). Amongst them were sites in genes such as *speC,* encoding an ornithine decarboxylase, *dgoT*, encoding a D-galactonate transporter and *ghrB*, encoding a glyoxylate/hydroxypyruvate reductase.

Outcome of the coselection and epistasis analysis was further visualised and investigated by using GWES-Explorer (https://github.com/jurikuronen/GWES-Explorer) (Supp. Fig. 3). Within the short-range links, ST131-B presented apparent loss events of many of the phage-related sites, including e.g. prophage host-nuclease inhibitor and phage tail proteins, which were maintained in ST131-B0 and ST131-C clades. Most ST131-B still harbouring the sites also showed an allele change compared to the loci in ST131-B0 and ST131-C. Oppositely, a distinct subclade in ST131-B showed a shared allele in phage recombination protein Bet to those of the other ST131 clades. As for the long-range links, most sites had shared alleles between ST131-B0 and ST131-C clades, differentiating them from ST131-B. An exception was the network of variants between leucyl aminopeptidase PepA and iron-sulfur stress protein YtfE and YtfB, where shared alleles differentiated most ST131-B0 and ST131-B isolates from the ST131-C clades.

### Plasmidome analyses

Altogether, 32 contigs of the OUCRU ST131-B0 hybrid assemblies were predicted as plasmid-derived. In addition, 578 plasmid-derived contigs were retrieved from the NORM collection (Arredondo-Alonso et al. 2025). These contigs were assigned to 15 different plasmid types (pT) (Fig. 3, Supp. Table 3), according to the existing mge-cluster *E. coli* plasmid typing scheme (Arredondo-Alonso et al. 2023; Arredondo-Alonso et al. 2025). While most pTs were disseminated across the phylogeny, some plasmid-clade association was seen in particular pTs (Supp. Table 3). Most of the plasmid contigs of pT2 (84.6%, 104/123), pT4 (77.6%, 59/76) and pT8 (95.0%, 76/80) were found in ST131-C1 and -C2, while pT5 included exclusively ST131-B plasmids. Also pT11 were harboured almost solely by ST131-B, with only 1 of 31 plasmids being in ST131-C1. pT10 corresponded to a singleton type with one ST131-B plasmid. Altogether seven of the pTs were harboured by OUCRU ST131-B0 isolates (pT0, pT2, pT3, pT4, pT7, pT8 and pT15). The remaining five types (pT1, pT12, pT13, pT14, pT16) were a mixture of ST131-B and ST131-C plasmids from the NORM collection (Fig. 3). In total, 110 plasmid-derived contigs remained unassigned to any plasmid type (pT-1) (Supp. Table 3).

**Figure 3.**
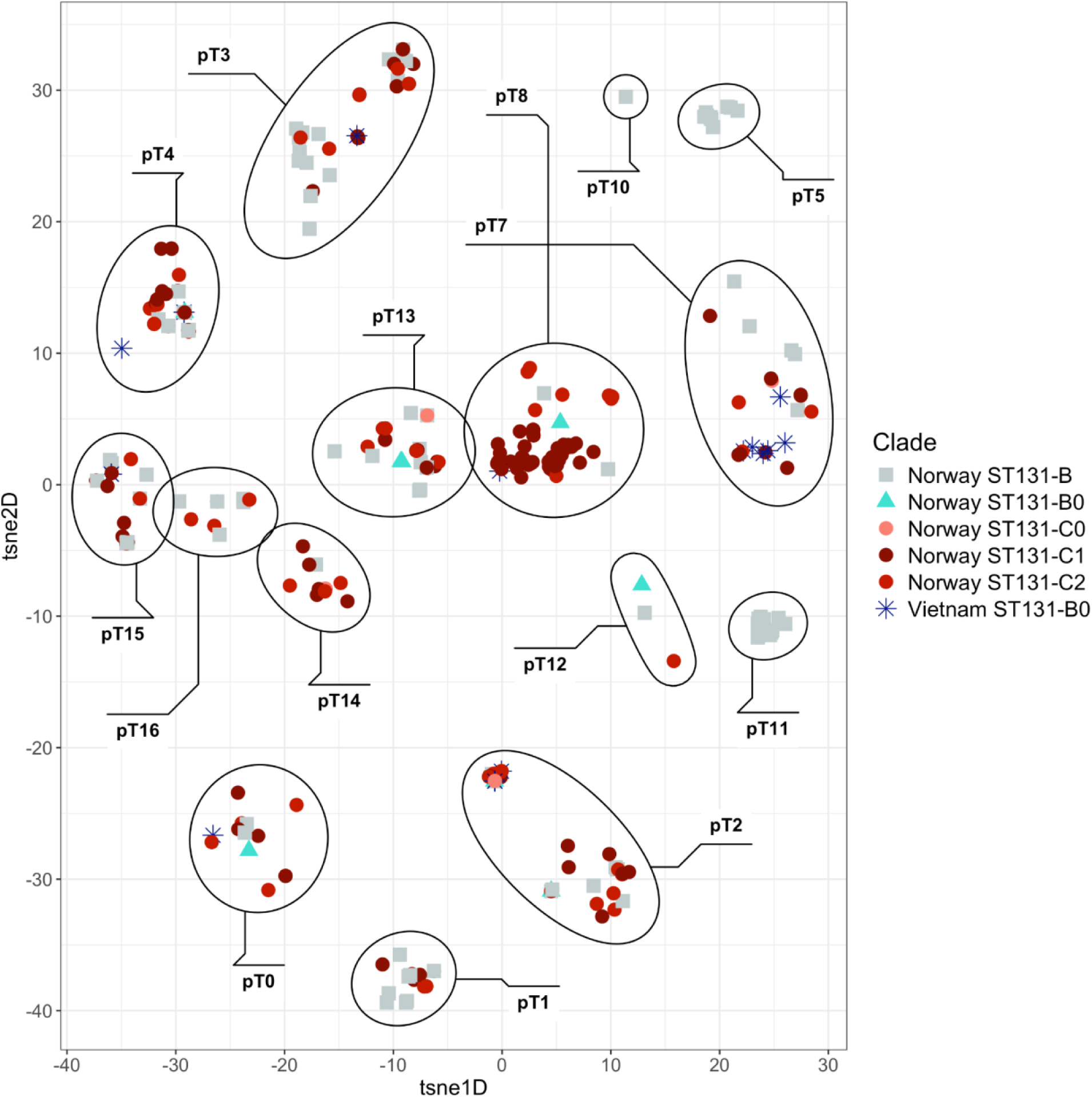
OpenTSNE embedding of 500 *E. coli* plasmid-predicted contigs of the Vietnamese ST131-B0 and Norwegian ST131-B, -B0, -C0, -C1 and -C2. Each point corresponds to a plasmid sequence and their assigned cluster is labelled by the plasmid type (n=15) defined by using mge-cluster with the existing cluster scheme (perplexity=62) (Arredondo-Alonso et al. 2025). Sequences are coloured and shaped by the country of origin and ST131 clade as indicated in the legend. The ellipses (in black) delimit the cluster coordinates. Plasmid-predicted contigs unassigned to any plasmid type (pT-1) were removed.

Of all the pTs, pT7 was found in most OUCRU ST131-B0 plasmid-predicted contigs in total (n=6), while pT2 was found in most OUCRU isolates (n=5) (Figs 1 and 3). Plasmids within the pT7 shared an IncFII replicon (Fig. 4A) and were average in size with most variation of all the pTs (mean 73.9 kbp, SD 35.1 kbp). Only two pT7 plasmids harboured CTX-M type β-lactamases: CTX-M-15 in an OUCRU ST131-B0 and CTX-M-19 in a NORM ST131-C1 plasmid (Fig. 4A), whereas six pT7 plasmids harboured *bla*_TEM-1_ β-lactamase genes. Of these, the OUCRU ST131-B0 plasmid carried both CTX-M-15 and TEM-1, notably colocalised with type II toxin-antitoxin systems. The synteny analyses within pT7 further depicted the loss of multiple colicin loci in the path from ST131-B to the ST131-C clades, which had already disappeared in the ST131-B0 (Fig. 4A). Of note, none of the ST131-B0 plasmids harboured microcin (colicin V) that was found widely present in ST131-B plasmids, particularly in pT5 (Supp. Table 3). The pTs corresponding to the largest subsets of plasmids in total were pT2 (n=123) and pT8 (n=80), the latter of which corresponds to the previously described plasmid type with high prevalence in ST131-C1 and frequently harbouring β-lactamase genes (Arredondo-Alonso et al. 2025). The pT8 plasmids also shared an IncFII replicon type (Fig. 4B), were amongst the largest in length (mean 115.9 kbp, SD 25.4 kbp) and 37.5% (30/80) harboured CTX-M-14, −15 or −27. pT2 plasmids, on the contrary, were amongst the smallest ones (mean 4.9 kbp, SD 0.8 kbp) and harboured no known AMR genes.

**Figure 4.**
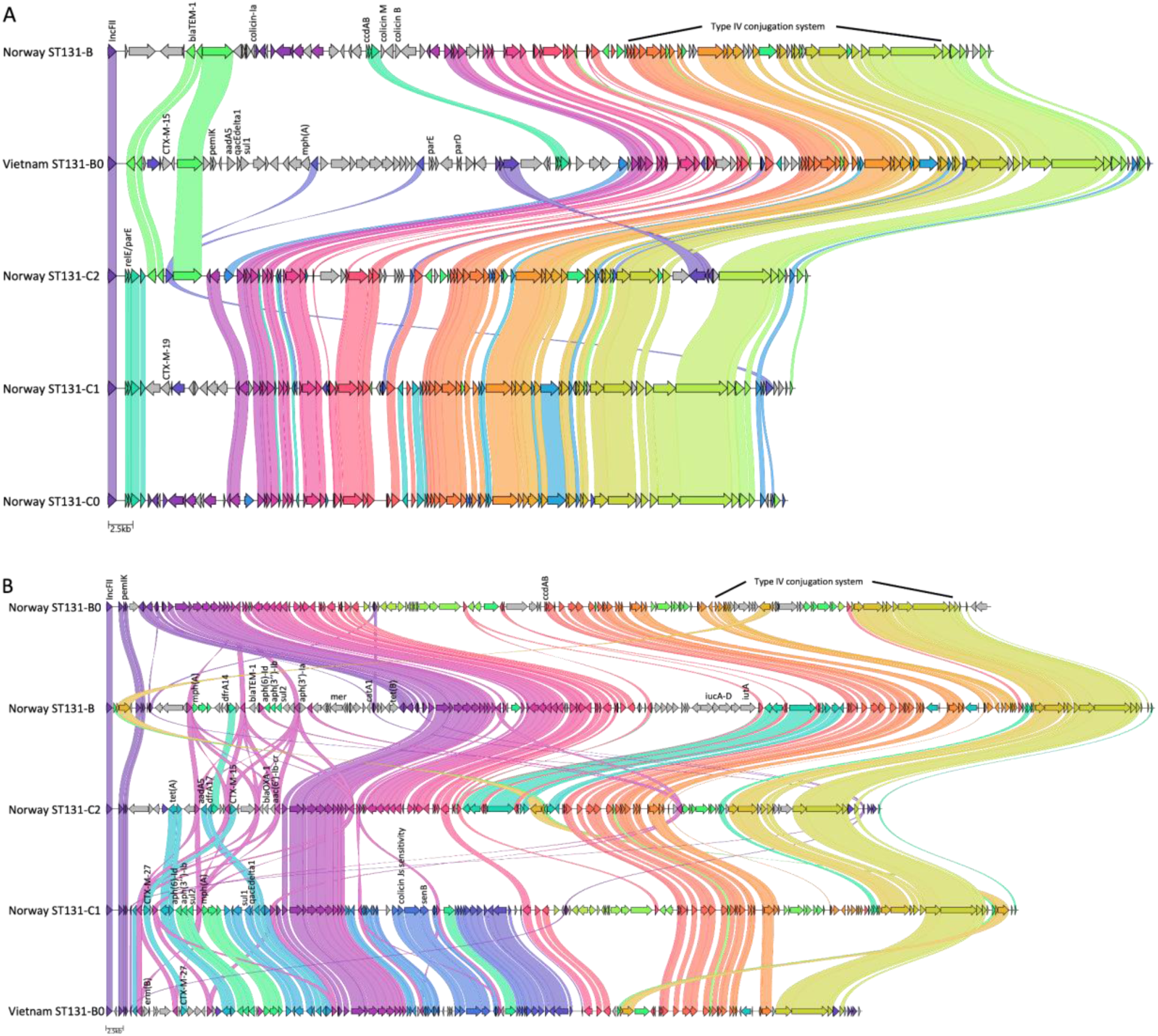
Synteny analysis of representative plasmid-predicted contigs of each ST131 clade within pT7 (A) and pT8 (B) plasmid clusters aligned by using Clinker (Gilchrist and Chooi 2021) with 80% sequence identity threshold. AMR, virulence and stress resistance genes were annotated using AMRFinderPlus (Feldgarden et al. 2021). Replication initiation proteins, conjugation machinery, colicins and type II toxin-antitoxin systems were retrieved from Bakta annotations (Schwengers et al. 2021). Within shared sequences, only the topmost annotations are noted.

To investigate potential co-selection of genomic variation within pT7 and pT8 plasmids that were evaluated to share evolutionary origin, the plasmid contigs were collectively analysed using the alignment-free PAN-GWES pipeline (Kuronen et al. 2024). Annotated identified links between variation included essential mobile genetic element functions under potential coselection, such as type IV conjugative transfer system coupling protein-encoding *traD* and type IV secretion system protein-encoding *traC* (Supp. Fig. 4). In addition, *traC* was found to pair with a partial overlap on a hypothetical protein-encoding *yhfA*.

## Discussion

The rapid rise of the *E. coli* ST131 lineage in clinical settings from the early 2000s onwards has inspired a considerable body of evolutionary research. The current consensus asserts that the success of the extantly circulating major ST131 subclades cannot be pinned down to a single genetic factor. Substantial evidence has accumulated to support the roles of metabolic and colonisation competition factors, as well as virulence and AMR determinants (McNally et al. 2019; Cummins et al. 2022; Arredondo-Alonso et al. 2025). Systematic population-based comparison of ST131 in asymptomatic colonisation versus bloodstream infections has further revealed that the ST131-A and ST131-B clades are relatively benign compared with ST131-C1 and ST131-C2 (Mäklin et al. 2022; Pöntinen et al. 2024), which are over-represented among infection cases. Further, recent study quantified that this is not primarily driven by any specific virulent capsule types, but corresponds instead to a lineage effect stemming from a larger set of genetic factors (Gladstone et al. 2025).

ST131-B is largely characterised as susceptible to antimicrobials compared to the ST131-C clades (Gladstone et al. 2021; Pöntinen et al. 2024; Arredondo-Alonso et al. 2025), although has in some cases been found to carry AMR traits (Reid et al. 2019; Roer et al. 2019; Saidenberg et al. 2020). ST131-B has been shown to be zoonotic, associated in particular with poultry meat (Liu et al. 2018), while in addition adapted to asymptomatic circulation among humans (Mäklin et al. 2022). The loss of microcin (colicin V) from the plasmids we observed here during evolutionary transition from ST131-B to ST131-B0 is well aligned with a restriction of the primary colonising host range to humans, since this microcin has been shown to be an important fitness determinant against Salmonella (Baquero et al. 2019). Expectedly, ST131-B would frequently compete with Salmonella when colonizing the gut of poultry. ST131-B0 has in turn been rarely observed in ST131 studies to date, but similarly to ST131-B, identified isolates are mainly from high-income countries and predominantly susceptible to antimicrobials. Overall, overrepresentation in high-income countries would be expected with more genomes sequenced. However, sequenced ST131-B0 isolates from China and Thailand were also found to lack *bla*_CTX-M_ genes (McNally et al. 2016; Stoesser et al. 2016; Gladstone et al. 2021).

The current consensus is that the ST131-C clades evolved from ST131-B through a *fimH* switch and by acquisition of mutational and plasmid-borne resistance traits (Ben Zakour et al. 2016; Matsumura et al. 2016; Pitout and Finn 2020). However, the lack of intermediary ST131-B0 isolates with complete genomic data, and particularly those CTX-M-positive, has resulted in a notable knowledge gap considering the more detailed evolutionary pathway of ST131 into MDR. In this study, combining high-quality sequence data and hybrid assemblies of clinical ST131-B0 isolates with a high-resolution *E. coli* ST131 cohort (Gladstone et al. 2021; Arredondo-Alonso et al. 2025), we were able to more thoroughly map the evolutionary path from ST131-B to ST131-C. Dating analyses of the combined cohorts placed ST131-B0 as an intermediary progenitor of ST131-C clades, as has been previously suggested (Ben Zakour et al. 2016; Pitout and DeVinney 2017). Moreover, our analyses presented a potential intermediary switch to and from the most common type *fimH54* in ST131-B0. For comparison, no ST131 isolates in the NORM cohort carried the *fimH54* allele, while in the entire Norwegian BSI cohort (n=3254) by (Gladstone et al. 2021), *fimH54* was detected in 141 isolates, majority of them in ST393 (26.2%), ST10 (21.3%) and ST95 (19.9%). Furthermore, most of the ST131-B0 isolates carried the ancestral K5 capsule type, preceding the higher capsule diversity seen in the ST131-C clades (Gladstone et al. 2025), and thereby further supporting their position as an intermediary ST131 clade.

In the combined collection of ST131 isolates from Vietnam and Norway, we found CTX-M-15 and CTX-M-27 to be the most common types of the CTX-M β-lactamases. We detected CTX-M-27 predominantly on plasmids while CTX-M-15 resided equally on plasmids and chromosomes, even with multiple copies on the same chromosome. Indeed, CTX-M-15 has been found in varying genetic contexts both on plasmids and chromosomes, particularly within *E. coli* ST131 (Lipworth et al. 2024). Of note, resistance levels conferred by β-lactamases are shown to be gene dosage-dependent (San Millan et al. 2016; Rodríguez-Beltrán et al. 2021). Thereby, carrying these genes on plasmids, which despite being relatively large in size can be harboured in multiple copies in a subset of cells in the colony, could provide a higher level of resistance to the bacteria regarded as a within-host community (Rodríguez-Beltrán et al. 2021). In the same line, upon chromosomal integration, like in the case of CTX-M-15 here, multiple copies may be beneficial for the pathogen due to gene dosage effects. Current evidence also shows AMR-carrying plasmids to be more plastic than others (Coluzzi and Rocha 2025) and recombination in general to be more frequent in genes carried on plasmids than those on chromosomes (Rodríguez-Beltrán et al. 2015), further suggesting that plasmid-borne genes evolve at a faster rate than chromosomal ones. That could also support the mutational evolution of CTX-M-27 from the CTX-M-14 background, possibly resulting in its position as one of the dominant alleles in our data.

Notably, we found all of the Vietnamese ST131-B0 isolates harboured CTX-M-type ESBLs, a trait previously thought of as distinctive for ST131-C clades. Indeed, none of the Norwegian ST131-B0 nor ST131-B clade had any *bla*_CTX-M_ genes (Fig. 2). The CTX-M types in the Vietnamese ST131-B0 isolates were either CTX-M-15 or CTX-M-27 which have been most commonly seen in ST131-C2 and ST131-C1, respectively (Johnson et al. 2016; Matsumura et al. 2016; Gladstone et al. 2021). Furthermore, bar one chromosomal CTX-M-15 with 98.63% sequence identity, both types were each found with fully identical sequences on chromosomes as well as on plasmids in the Vietnamese ST131-B0 (Fig. 2). This indicates that, in addition to being readily mobile in ST131-B0, CTX-M-type β-lactamases may have already been established in the ST131 lineage prior to the branching event ultimately leading to the ST131-C clades, unless these were acquired in a separate event later.

We also found *bla*_CTX-M_ genes co-localising on plasmids with toxin-antitoxin systems (Fig. 4), possibly aiding in the distribution of AMR traits in the ST131-B0. Toxin-antitoxin systems are important mechanisms in plasmid stability and maintenance, and co-localisation of these systems together with AMR traits on conjugative plasmids are of concern due to the potential for dissemination and fixation of the resistance determinants in the bacterial populations. Of the type II TA systems, the ParDE super family has been identified as distributed in IncF-type plasmids in *E. coli*, also contributing to the stress tolerance of the pathogen (Kamruzzaman and Iredell 2019). This system was also present in the CTX-M-15-carrying IncF-type plasmid in a Vietnamese ST131-B0 isolate. While ParDE was absent in the Vietnamese CTX-M-27 plasmid, another two TA systems common in Enterobacteriaceae, CcdAB and PemIK (Wu et al. 2020), were found on ST131-B0 plasmids harbouring both CTX-M-15 and CTX-M-27.

Regarding unique genes found exclusively in ST131-B0 while absent in ST131-C clades, prevalence of the arginine deiminase (ADI) pathway (*arc* operon) has been shown to be higher in ESBL-producing *E. coli*, in particular those carrying CTX-M-type β-lactamases (Billard-Pomares et al. 2019). However, the study found it more common in IncF plasmids than chromosomes. As per *in vivo* competition experiments in a mouse model, it was suggested to be of advantage for the strain in UTI, while being costly in intestinal colonization. Our findings of gene gain and loss in the stepwise evolution towards ST131-C clades thus highlight further targets for experimental work to elucidate their underlying exact functional drivers. Similarly, recent evidence supports that the ST131-C clade genomes continue to undergo rapid diversification (Gladstone et al. 2025; Molari et al. 2025), and further research is needed to elucidate how much of this represents drift versus selection.

## Materials and Methods

### Genomic datasets

In total, 9 *E. coli* human clinical isolates from blood stream infections (BSI; n = 7) and urinary tract infections (UTI; n = 2) from Oxford University Clinical Research Unit (OUCRU), Ho Chi Minh City, Vietnam, collected in 2017 and 2018, were sequenced within this study by short and long-read techniques. In order to use data for comparison with complete genomes and both short- and long-read sequences available, a collection of BSI isolates from the Norwegian surveillance programme on resistant microbes (NORM) study, collected 2002–17 in Norway (Gladstone et al. 2021), was selected for comparative genomics analyses. Whole-genome sequences, short-read assemblies, *fimH* types and clade designations of 203 *E. coli* clonal complex (CC) 131 clades, here referred to as ST131-B, -B0, -C0, -C1 and -C2, were included in the analyses. For comparative pangenome and plasmidome analyses, available hybrid assemblies (n=197) and plasmid-predicted contigs (n=578) of the NORM isolates were retrieved from (Arredondo-Alonso et al. 2025).

For short-read sequencing of the OUCRU collection, whole-genome DNA was extracted using the Wizard Genomic DNA Purification Kit (Promega, Madison, USA) according to the manufacturer’s instructions and sequencing was performed at the Wellcome Sanger Institute, UK, using Illumina HiSeq X Ten paired end platform. Mixed strains and species contamination were excluded by using Kraken v.0.10.6 (Wood and Salzberg 2014). Sequencing quality was controlled by 30x minimum depth of coverage, and maximum of 100 contigs, 0.20 highest minor allele frequency in multi-locus sequence type (MLST) genes and 500 heterozygous SNP sites. Short-read sequence data was assembled by using a published pipeline with Velvet optimisation (Page et al. 2016) to coincide with the NORM dataset.

For long-read sequencing on the Oxford Nanopore Technologies (ONT) platform, whole-genome DNA of the OUCRU isolates was extracted using an in house phenol:chloroform extraction procedure. Concentration and DNA integrity were measured using NanoDropOne spectrophotometer (Thermo Fisher Scientific, Waltham, USA) and the Qubit dsDNA HS assay kit (Thermo Fisher Scientific) on a CLARIOstar microplate reader (BMG Labtech, Ortenberg, Germany). The ONT libraries were prepared using SQK-NBD112-24 barcoding kit for 24-barcoding runs, on FLO-MIN106 flow cells. Sequencing was run for 72 hours on GridION (Oxford Nanopore Technologies, Oxford, UK) using MinKNOW Core v.5.8.6. Basecalling and demultiplexing were performed using Guppy v.6.4.6 with high-accuracy basecalling. ONT long reads were combined with the short-read data using Hybracter v.0.10.0 with default parameters (read length >1 kbp and read quality score of >9 for long reads) and recommended estimated genome size of 4 Mbp for *E. coli* (Bouras et al. 2024), resulting in 6 complete and 3 incomplete hybrid assemblies.

### Sequence typing

ST131 sequence type of the OUCRU isolates was confirmed by using SRST2 v.0.2.0 (https://github.com/katholt/srst2) by comparing short-read data to the *Escherichia* species MLST database (https://pubmlst.org/organisms/escherichia-spp, retrieved 9th May 2025). *fimH* typing was performed on short-read assemblies by using Abricate v.1.0.1 (https://github.com/tseemann/abricate) on the fimtyper database (https://bitbucket.org/genomicepidemiology/fimtyper_db/downloads/, retrieved 9th May 2025), and ST131 clade B0 was further assigned by clade-specific SNPs (Price et al. 2013; Ben Zakour et al. 2016) and phylogenetic comparison to the NORM ST131.

*E. coli* O-H serotypes of both the OUCRU and NORM isolates were assigned on short-read assemblies (n=212) by using Abricate v.1.0.1 (https://github.com/tseemann/abricate) on the EcOH database (Ingle et al. 2016). Capsule typing of both collections was performed on the short-read assemblies (n=212) by using Kaptive v.3.1.0 (Stanton et al. 2025) against the *E. coli* group 2 and 3 capsule database v.2.0.0 (Gladstone et al. 2025). Only typeable loci without any identified problems were accepted.

### *Bla*_CTX-M_ screening

To investigate presence of and differentiating chromosomal *vs* plasmid loci and number of chromosomal copies of the *bla*_CTX-M_-type ESBL genes in the OUCRU and NORM isolates, the resistance genes were screened from the hybrid assemblies (n=206) using AMRFinderPlus v.4.0.22 (Feldgarden et al. 2021) with database v.2025-03-25.1 and *Escherichia* taxonomy group (--organism Escherichia). Minimum identity was set to 75% and minimum coverage to 80%.

### Dating analyses

For dating analyses of the combined OUCRU and NORM isolates (n=212), short reads were mapped against the *E. coli* ST131 reference genome EC958 (HG941718.1) using Snippy v.4.6.0 (https://github.com/tseemann/snippy) and recombination was removed by using Gubbins v.3.4.0 (Croucher et al. 2015). Dated tree was inferred with BactDating v.1.1.1 (Didelot et al. 2018) in 3 separate runs, using the additive relaxed clock (ARC) model, 100,100,100 MCMC chain length and 100,000 thin, and verified for estimated sample size of over 200. The model fit was compared to that of random dates for significance of the temporal signal.

### Comparative genomics

For comparison of the pangenomes as well as the gene gain and loss rates between the ST131 clades in the combined collection, the hybrid assemblies (n=206) were annotated by using Bakta v.1.10.3 with the database v.5.0 and *Escherichia* genus (Schwengers et al. 2021). Pangenomes were estimated by using Panaroo v.1.5.2 (Tonkin-Hill et al. 2020) with sensitive mode, by removing invalid genes and merging paralogs, and with a minimum pairwise sequence-identity threshold of 70%. Gene gain and loss rates between ST131-B0, ST131-B, ST131-C1 and ST131-C2 were estimated and compared using Panstripe v.0.3.3 assuming a Gaussian distribution (Tonkin-Hill et al. 2023). The combined dated phylogeny was used to partition ST131-C1 and ST131-C2 for the analysis, while a separately dated phylogeny was used in partitioning ST131-B and ST131-B0.

Potential coselection and pangenome-spanning epistasis of genomic variation was investigated from multiple-sequence alignment against the *E. coli* ST131 reference genome EC958 (HG941718.1) by using LDWeaver v.1.5.2 (Mallawaarachchi et al. 2024) and further visually analysed by using GWES-Explorer (https://github.com/jurikuronen/GWES-Explorer). EC958 was annotated using Bakta v.1.9.3 with the database v.5.1 and *Escherichia* genus (Schwengers et al. 2021) and used in mapping and annotation of the top hits.

### Plasmidome analyses

For comparative plasmidome analyses of the combined OUCRU and NORM datasets, plasmid prediction of the OUCRU collection was performed to coincide with the NORM analyses (Arredondo-Alonso et al. 2025): Hybrid assembly contigs smaller than 1 kbp and larger than 300 kbp were filtered out. Plasmid contigs were then predicted by using mlplasmids v.2.1.0 with *Escherichia coli* species model and minimum plasmid probability threshold of 0.3 (Arredondo-Alonso et al. 2018) and by using geNomad v.1.7.0 end-to-end command with minimum plasmid score of 0.7 (Camargo et al. 2024). Remaining contigs were classified as plasmid-predicted if they contained a replication type as defined by mob-typer v.3.1.4 (Robertson and Nash 2018; Robertson et al. 2020). Predicted plasmid contigs were then delineated into plasmid types (pT) by using mge-cluster v.1.1.0 (Arredondo-Alonso et al. 2023) against the existing *E. coli* scheme with a perplexity value of 62 (Arredondo-Alonso et al. 2025).

Representative plasmids of each available ST131 clade, opting for ESBL β-lactamases when present, were selected for within-pT comparison in pT7 and pT8. The starting position of the sequences was rotated to IncFII by using Circlator v.1.5.5 (Hunt et al. 2015) with the fixstart command. The plasmids were then screened for AMR, virulence and stress resistance genes using AMRFinderPlus v.4.0.22 (Feldgarden et al. 2021) with database v.2025-03-25.1 and *Escherichia* taxonomy group (--organism Escherichia), with minimum identity of 75% and minimum coverage of 80%. The plasmids were further annotated using Bakta v.1.7.0 with the database v.5.0 and *Escherichia* genus (Schwengers et al. 2021) and gene synteny analysis performed using Clinker v.0.0.31 (Gilchrist and Chooi 2021) with minimum alignment sequence identity of 80%.

Potential coselection and epistasis of genomic variation within pT7 and pT8 plasmids was investigated directly from predicted plasmid contigs (n=103) by using phenotype- and alignment-free PAN-GWES pipeline (Kuronen et al. 2024), implementing cuttlefish v.2.2.0 (Khan and Patro 2021; Khan et al. 2022) and SpydrPick v.1.2.0 (Puranen et al. 2018; Pensar et al. 2019). K-mer length for PAN-GWES was set to 81 bp that was estimated to capture fine-scale mutations, while allowing for enough sequence diversity and capturing differences at gene level. Unitig pairs with an average distance exceeding the standard deviation of their distances and pairs present in less than 5% of the plasmid contigs were filtered out. Selected outlier top hits were annotated using annotate_hits_pyseer of the pyseer tool v.1.3.12 (Lees et al. 2018) against selected reference plasmid (28328_1#6_plasmid00001; 106718 bp) that was confirmed to cover for all of the hits.

All visualisations in the study were performed in R v.4.5.0 or higher (R Core Team 2020), Clinker v.0.0.31 (Gilchrist and Chooi 2021) and Microreact (Argimón et al. 2016).

## Data availability

Short-read Illumina and long-read ONT sequence reads generated within this study are publicly available at the European Nucleotide Archive (ENA) under the study ID PRJEB29743 and PRJNA1370503, respectively. The Norwegian short-read data is publicly available at the ENA under the study ID PRJEB32059 and long-read data under the study IDs PRJEB45354 and PRJEB57633. Descriptive data on the isolates together with a dated phylogeny is publicly available as a Microreact project [https://microreact.org/project/ecoli-st131-evolution].

## Acknowledgements

We acknowledge the support from the Wellcome Sanger Institute sequencing facility and the Genomics Support Center Tromsø.

## Author contributions

Conceptualisation - AKP, NTD, NVVC, NTH, JC

Funding acquisition and resources - NVVC, JP, NTH, JC,

Methodology, formal analysis - AKP, NVT, SM, JK, HXY, PLKY, NPHL, JC

Validation - AKP, SM, RAG, ØS, GT-H, JC

Visualisation - AKP, SM

Writing, original draft - AKP with contributions from JC

Writing, review and editing - all authors

