## Supplemental Figures for "Mapping the evolutionary path towards multi-drug resistance in the pandemic *Escherichia coli* ST131 lineage"

Affiliations
1 Department of Biostatistics, Faculty of Medicine, University of Oslo, Oslo, Norway
2 Oxford University Clinical Research Unit, Ho Chi Minh City, Vietnam
3 Faculty of Veterinary Medicine, College of Agriculture, Can Tho University, Can Tho, Vietnam
4 Peter MacCallum Cancer Centre, Melbourne, Australia
5 Sir Peter MacCallum Department of Oncology, The University of Melbourne, Melbourne, Australia
6 Norwegian Centre for Detection of Antimicrobial Resistance, Department of Microbiology and Infection Control, University Hospital of North Norway, Tromsø, Norway
7 Hospital for Tropical Diseases, Ho Chi Minh City, Vietnam
8 Department of Health, Ho Chi Minh City, Vietnam
9 Department of Veterinary Medicine, University of Cambridge, Cambridge, United Kingdom
10 Department of Microbiology and Immunology, The University of Melbourne at the Peter Doherty Institute for Infection and Immunity, Melbourne, Australia
11 Nuffield Department of Medicine, Centre for Tropical Medicine, University of Oxford, Oxford, United Kingdom
12 Center for BioMedicine Research, Pham Ngoc Thach University of Medicine, Ho Chi Minh City, Vietnam
13 Parasites and Microbes, Wellcome Sanger Institute, Cambridge, United Kingdom
14 Helsinki Institute of Information Technology, Department of Mathematics and Statistics, University of Helsinki, Helsinki, Finland

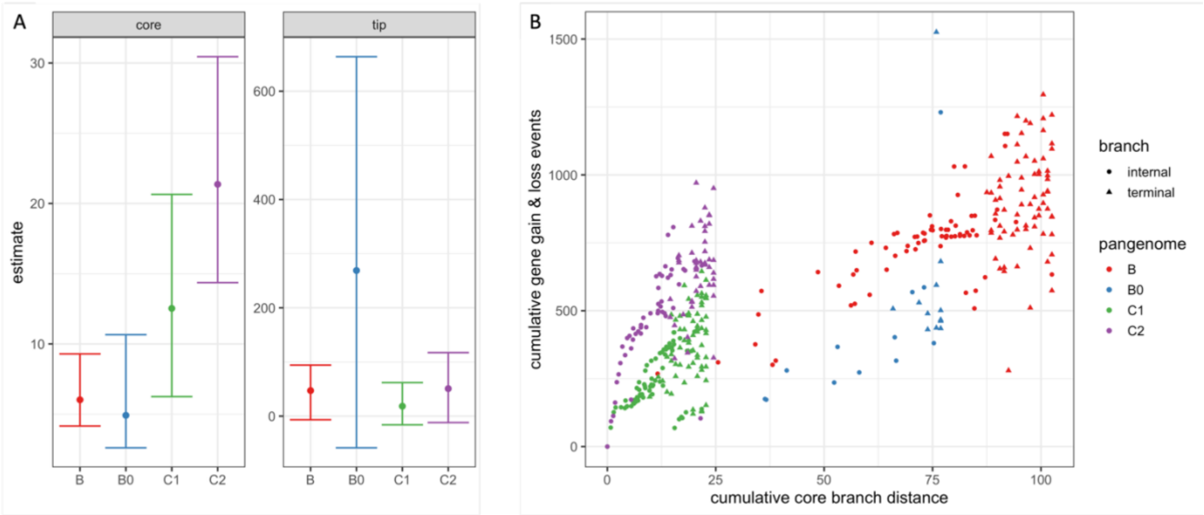


**Fig. S1. Comparison of the gene gain and loss rates in *E. coli* ST131-B, ST131-B0, ST131-C1 and ST131-C2 using Panstripe (Tonkin-Hill *et al.* 2023).** (A) The estimated parameters of the generalized linear model. Higher values of the core estimates indicate increased gene gain and loss rates, while the tip estimates describe gene gain and loss events occurring only at the tips of the phylogenies. Error bars represent the 95% confidence interval of the parameter estimates. (B) The estimated cumulative gene gain and loss events in each clade.

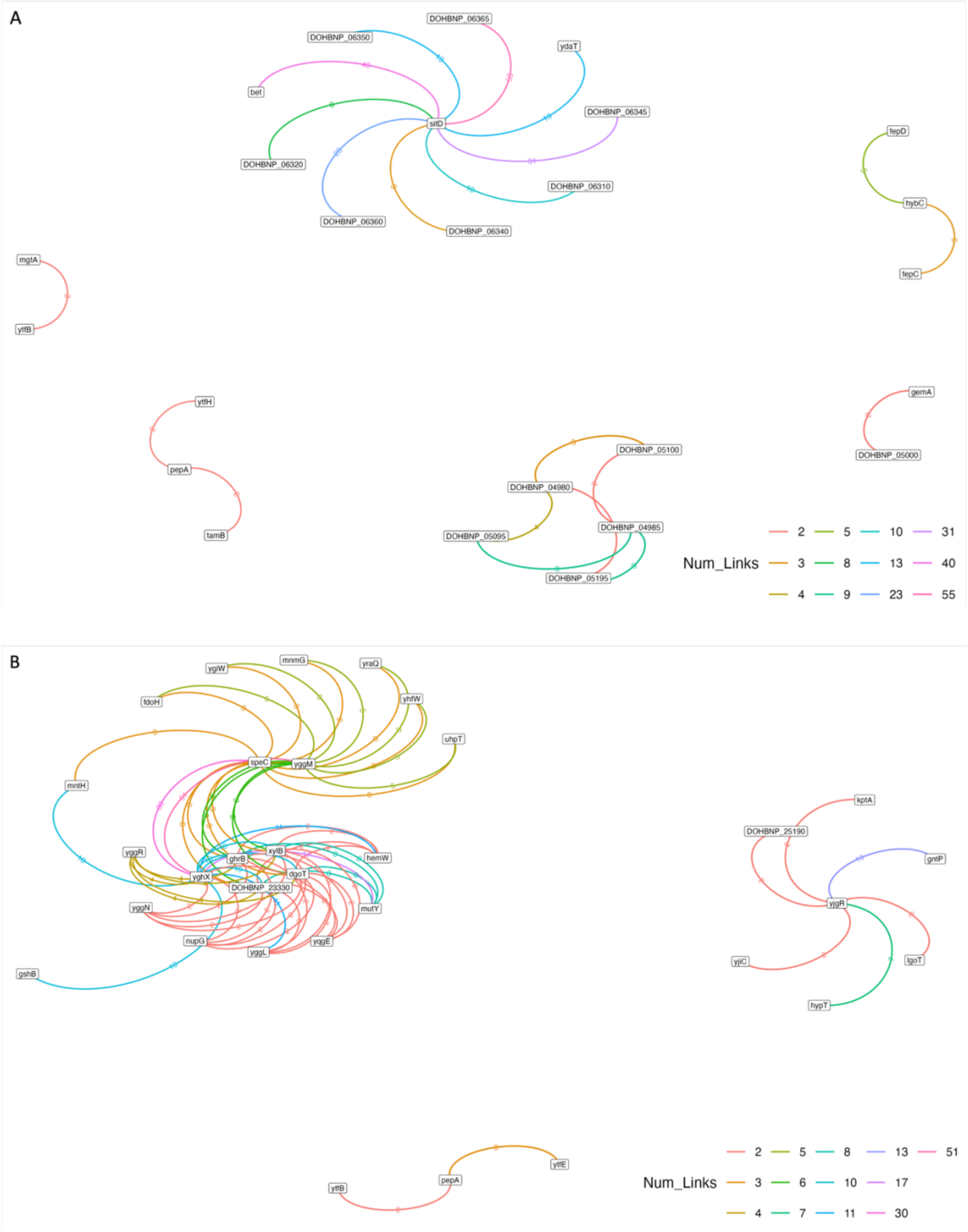


**Fig. S2. Investigation of potential co-selection and pangenome-spanning epistasis of genomic variation.** Networks in (A) short-range and (B) long-range highest scoring hits defined by LDWeaver (Mallawaarachchi *et al.* 2024). The edges are coloured according to the number of links between genomic region nodes as indicated in the legends. Annotations and SNP positions are based on the EC958 reference genome (HG941718.1).

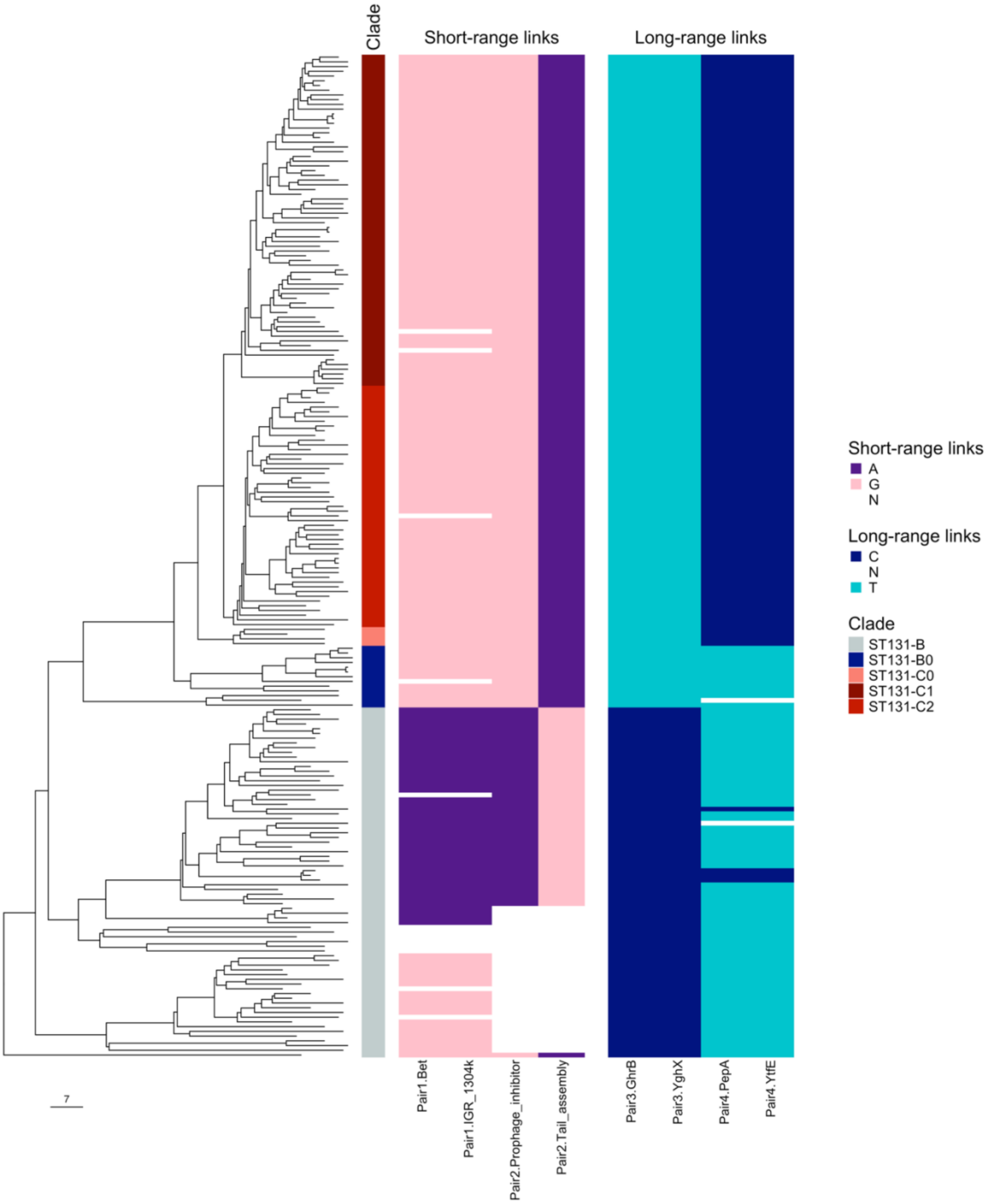


**Fig. S3. Investigation of allelic variation in potential coselection**. Variation in selected highest scoring loci for short- and long-range link pairs is shown by the heatmaps, aligned with the ST131 clades and the dated phylogeny where the tree scale indicates years. The key shows the colour for each nucleotide, with N indicating a missing loci. Protein annotations are based on the EC958 reference genome (HG941718.1). IGR = intergenic region.

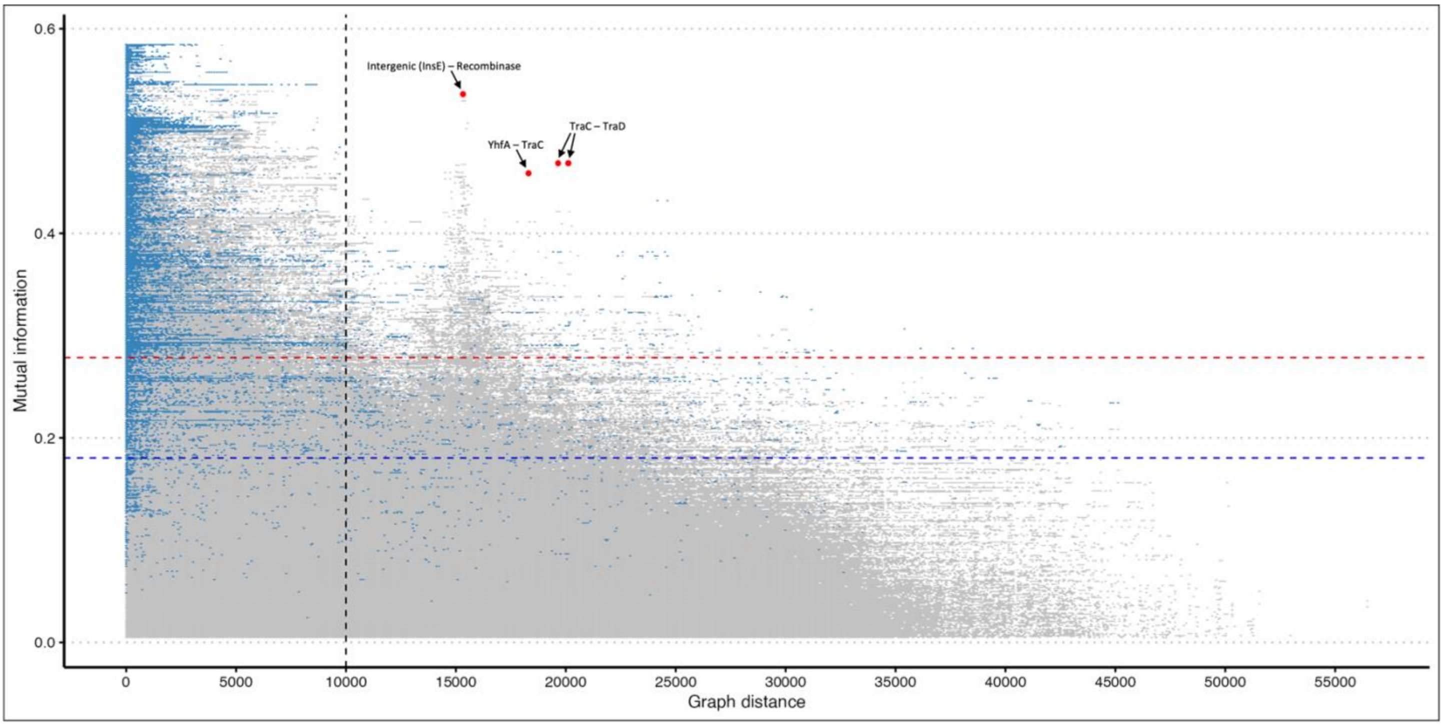


**Fig. S4. Investigation of potential coselection and epistasis of genomic variation within predicted pT7 and pT8 plasmid contigs (n=103) by using PAN-GWES pipeline (Kuronen *et al.* 2024).** A Manhattan plot indicating the strength of linkage disequilibrium (LD) measured by using mutual information (MI) *vs* genomic distance in the graph (bp). Unitig pairs present in less than 5% of the predicted plasmid contigs and with an average distance greater than the standard deviation of their distances were filtered out. Selected strong links of potential coselection are indicated by red dots. These included links between a putative site-specific recombinase and an intergenic site upstream of plasmid transposase InsE-enconding gene and between a hypothetical protein-encoding *yhfA* and type IV secretion system protein TraC as well as TraC and type IV conjugative transfer system coupling protein TraD. Horizontal blue and red lines indicate the outlier and extreme outlier thresholds, respectively, and the vertical black line indicates the estimated background LD threshold below which it is difficult to differentiate outlier peaks.
